# Deep learning framework for kinematic event detection and stimulation decoding in primate reaching behavior

**DOI:** 10.64898/2026.07.14.738537

**Authors:** Avital Markus, Nirvik Sinha, Yifat Prut, Jacob Goldberger

**Author notes:** These authors contributed equally to this study (Avital Markus, Nirvik Sinha). These authors contributed equally to this study (Yifat Prut, Jacob Goldberger). Corresponding authors’ (J.G.), (Y.P.).

## Abstract

Accurate analysis of motor behavior requires the reliable detection of ongoing kinematic events and a granular characterization of the changes in motor output that occur in response to neural impairments. This article describes a deep learning framework based on bidirectional long short-term memory (BiLSTM) networks developed to analyze single-trial three-dimensional reaching trajectories recorded from two non-human primates. The framework was used to detect movement onset and corrective turning points, and to differentiate control from the perturbed trials during which cerebellar output was blocked. This approach was compared to manually annotated data. Only small within-animal errors in detecting movement onset times were observed (8.92 ± 2.03 ms and 9.56 ± 3.64 ms for the two monkeys). These errors were significantly smaller (p < 0.001) than those obtained using conventional velocity-threshold methods. Transferring the same detection algorithm from one animal to the other resulted in large errors because the reconstructed workspaces were represented in different coordinate frames. Orthogonal Procrustes alignment substantially reduced the between-animal event-detection errors, and brought performance closer to the within-animal range. Decoding of the perturbed vs. the control trials achieved above-chance levels of accuracy for each animal (accuracies of 71.0% and 61.9% respectively). However, the between-animal generalization was poor (near chance level) and was not improved by geometric alignment. These findings suggest that geometric alignment can support the transfer of shared kinematic event structure between animals, but that perturbation-related changes in movements reflect animal-specific compensatory strategies which cannot be generalized.

## Introduction

Motor behavior provides a sensitive reflection of the state of the nervous system. In fact, several neurological disorders first manifest as abnormalities in motor output. Therefore precise and robust quantification of motor behavior is essential for studying both the physiology and the pathophysiology of the nervous system [1, 2]. Meaningful assessment of motor behavior typically requires movement to be performed under natural, unconstrained conditions, yet this very requirement complicates rigorous and reproducible behavioral quantification [3]. To address this challenge, recent years have seen the development of automated tools for accurate tracking of unconstrained behavior [4–6], complemented by algorithms that segment continuous behavior into discrete behavioral epochs [7, 8]. Most of these approaches, however, were designed for relatively stereotyped rodent behaviors such as locomotion. In contrast, motor behavior in humans and non-human primates is substantially more complex and variable, both within and across individuals, necessitating dedicated solutions [7, 8]. In primates motor neuroscience, experimental paradigms frequently focus on upper-limb movements, where quantitative behavioral analysis requires millisecond-precision detection of kinematic landmarks such as movement onset and corrective turning points [2]. These points serve as critical temporal anchors for aligning neural activity with behavior and provide sensitive, non-invasive markers of motor dysfunction associated with neurological disease [1, 2].

To-date, extracting reliable and generalizable kinematic events from noisy, variable, and subject-specific movement trajectories remains a major computational challenge. Most existing approaches identify movement events using heuristic criteria, such as fixed velocity thresholds (e.g., 15% of peak velocity), that require manual parameter tuning and are therefore sensitive to noise, baseline drift, and threshold selection [9–12]. Furthermore, these parameters are often optimized independently for each subject to accommodate differences in motor behavior and recording conditions, limiting reproducibility and compromising quantitative comparisons across subjects. More principled statistical approaches have also been proposed, including log-likelihood ratio tests applied to whitened velocity signals [13] and adaptive autoregressive change-point detection algorithms [14]. Although these methods reduce reliance on heuristic thresholds, they typically require assumptions about signal statistics or model parameters that may not generalize across experimental conditions. Dynamic time warping (DTW) has likewise been used to align movement trajectories, but without appropriate constraints it can generate non-physiological temporal correspondences that complicate the identification of meaningful kinematic events [15].

To address the limitations of sensitivity and generalizability, we developed a deep learning framework based on bidirectional long short-term memory (BiLSTM) networks, which are designed to model long-range temporal dependencies through gated memory mechanisms while processing trajectories in both forward and backward temporal directions [16, 17]. The bidirectional architecture is well suited for kinematic sequence analysis because the timing of movement events depends not only on preceding motion but also on the subsequent evolution of the trajectory. By integrating information from both temporal contexts, the model can identify kinematic events more robustly across noisy and variable movement trajectories.

This proposed framework comprises three complementary components. First, it includes a BiLSTM-based event detection algorithm that identifies key kinematic events from movement trajectories. Second, it incorporates an inter-animal alignment procedure based on the orthogonal Procrustes algorithm [18], which compensates for arbitrary differences in coordinate frames across experimental setups and enables trained event detectors to generalize across animals Third, it implements a single-trial decoding approach that detects subtle changes in movement kinematics induced by experimental perturbations.

We validated the framework using markerless video recordings of two monkeys performing a center-out reaching task. On a trial-by-trial basis, the BiLSTM accurately detected both movement onset and the corrective turning point elicited by an unexpected displacement of the instructed target. Following geometric alignment, event detection generalized successfully across animals, demonstrating that the model can be transferred despite differences in recording geometry. Moreover, movement onset was identified with substantially lower mean absolute error than a conventional velocity threshold-based approach.

We next evaluated whether the same network architecture could detect perturbation-induced changes in movement kinematics at the single-trial level. Using trajectories recorded during high-frequency stimulation that transiently blocked cerebellar output, the network discriminated control and perturbed trials with higher accuracy than approaches based on endpoint kinematics or trial-averaged trajectories, highlighting the importance of preserving the temporal structure of movement dynamics. Although decoding performance was consistently above chance, accuracy varied across movement directions and did not generalize between animals, suggesting that subject-specific motor strategies contribute to perturbation-related changes that cannot be fully normalized by geometric alignment alone.

Together, these findings demonstrate that the proposed framework provides a robust and generalizable approach for detecting behaviorally meaningful kinematic events in unconstrained primate movements while remaining sufficiently sensitive to identify subtle, perturbation-induced changes in motor behavior at the single-trial level.

## Results

Two adult female macaques (*Macaca fascicularis*) were trained to perform an instructed-delay center-out reaching task on a vertical touchscreen (Fig. 1a,b). Each trial began with acquisition of a central target, followed by presentation of one of eight peripheral targets. After a variable delay period, disappearance of the central target served as the movement initiation cue, instructing the monkey to reach to the cued peripheral target to obtain a reward. In 20% of trials, the peripheral target unexpectedly switched to a different location shortly after the GO cue, requiring online updating of the planned or ongoing reach (hereafter, referred to as *update trials*).

**Figure 1.**
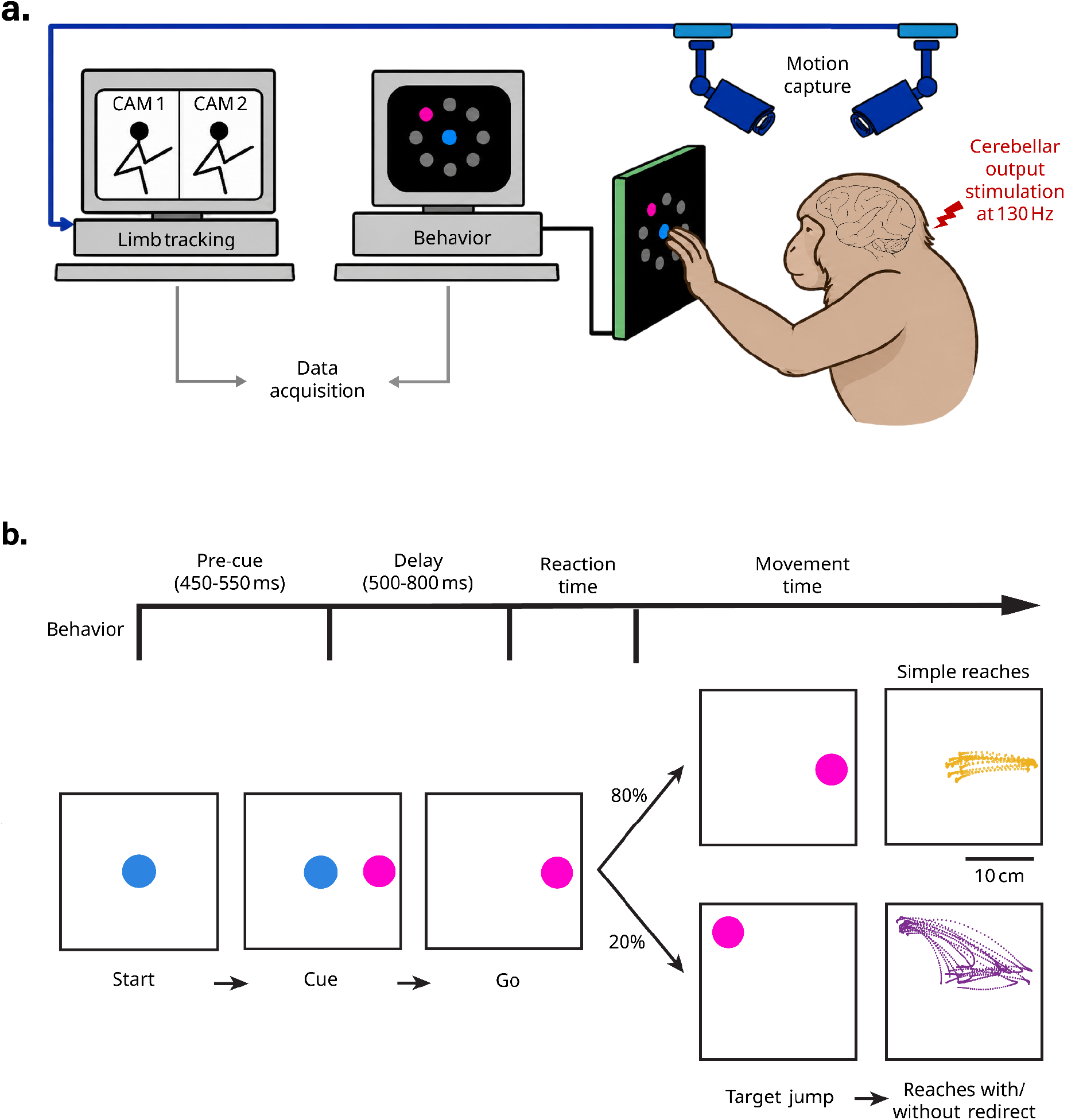
Experimental setup and behavioral paradigm. (a) Schematic of a monkey performing the delayed reaching task on the touchscreen. The behavior of the monkey was recorded using 120 Hz cameras synced online with the on-screen behavioral events. For a subset of trials, the cerebellar output to the motor cortex was disrupted by 130 Hz stimulation through a chronically implanted electrode in the superior cerebellar peduncle. (b) Sequence of screen events during the performance of a trial. Each trial began with the monkey maintaining touch within a central target. Subsequently, one of eight peripheral targets appeared (cue). After a delay, the central target disappeared (‘Go’ signal), prompting the monkey to reach for the peripheral target and briefly hold the cursor there to receive a reward. In 20% of the trials, the cued target jumped to a different location after a variable period from the ‘Go’ signal. Depending on jump timing relative to movement onset, the monkey either reached directly toward the final target or initiated a reach toward the first target and corrected online toward the final target.

Experiments consisted of alternating control and perturbation blocks of approximately 60 trials each. During perturbation blocks, cerebellar output to the motor cortex was reversibly disrupted by high-frequency stimulation (130 Hz) of the superior cerebellar peduncle delivered through a chronically implanted electrode [19]. This manipulation produces robust deficits in reaching movements that resemble key features of cerebellar ataxia [19,20].

The dataset comprised 13,330 trials from monkey T, including 1,974 update trials, and 10,204 trials from monkey N, including 656 update trials. These data were used for both kinematic event detection and single-trial decoding of control versus cerebellar perturbation conditions.

### Kinematic event detection within and across animals

We first assessed the ability of the BiLSTM model to accurately detect kinematic events from single-trial hand trajectories when both training and testing were performed on data from the same animal (i.e., within-animal training and testing). Model predictions were evaluated against manually annotated ground-truth events. Unless otherwise stated, all analyses were repeated over 10 independent training runs, and results are reported as mean ± standard deviation. The model identified movement onset with high temporal precision in both monkeys, yielding a mean absolute error (MAE) of 8.92 ± 2.03 ms for monkey T and 9.56 ± 3.64 ms for monkey N. Detection of trajectory turning points was likewise accurate, although less precise than movement onset, with MAEs of 23.93 ± 4.83 ms and 39.31 ± 5.66 ms for monkeys T and N, respectively. These findings demonstrate that the BiLSTM reliably captures the temporal relationship between hand kinematics and behaviorally defined events, enabling accurate event detection at the single-trial level.

We next compared the BiLSTM against a conventional heuristic approach for movement-onset detection based on hand kinematics [9,10]. As a baseline, movement onset was defined as the first time point at which hand velocity exceeded 20% of the trial-specific peak velocity. Across both animals, the BiLSTM substantially outperformed this threshold-based method. In monkey T, the heuristic approach yielded a mean absolute error (MAE) of 23.41 ms, compared with 8.92 ± 2.03 ms for the BiLSTM (Wilcoxon signed-rank test, *p* < 0.001). Similarly, in monkey N, the threshold-based method produced an MAE of 18.98 ms, whereas the BiLSTM achieved an MAE of 9.56 ± 3.64 ms (Wilcoxon signed-rank test, *p* < 0.001). These results demonstrate that the BiLSTM provides substantially more accurate and robust movement-onset detection than a widely used heuristic kinematic criterion.

We next asked whether a model trained on one animal could generalize directly to another without additional adaptation. Despite both monkeys performing the same task under identical experimental conditions, cross-animal generalization was poor. Training the model on monkey T and testing it on monkey N increased the mean absolute error (MAE) to 370.45 ± 34.40 ms for movement onset and 162.85 ± 27.78 ms for turning-point detection. Conversely, training on monkey N and testing on monkey T yielded MAEs of 278.14 ± 20.65 ms and 113.80 ± 27.16 ms for movement onset and turning-point detection, respectively. These substantial increases in error demonstrate that the learned kinematic representations were not directly transferable across animals, motivating the need for an alignment procedure to compensate for inter-subject differences.

The poor cross-animal performance was consistent with differences in the geometric organization of the recorded target spaces. Although both monkeys performed the same reaching task, the eight target-direction vectors were represented in differently oriented coordinate frames owing to the arbitrary world coordinates introduced by the three-dimensional reconstruction procedure (Fig. 2). To compensate for this mismatch, we aligned the datasets using an orthogonal Procrustes transformation [18], thereby bringing the target configurations into a common reference frame.

**Figure 2.**
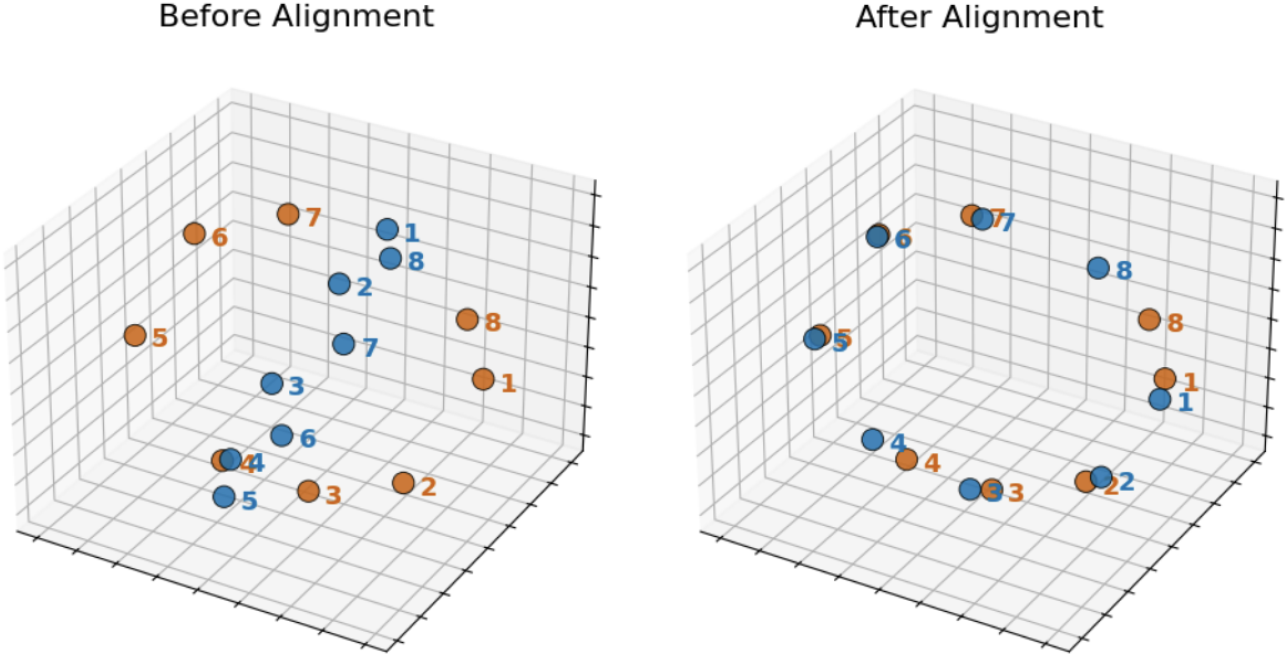
Visualization of the eight target vectors across monkeys. Left: target directions before alignment, showing inconsistent orientations between monkeys. Right: target directions after applying the IAA algorithm, bringing trajectories into a common reference frame. Colors indicate subjects: T (orange) and N (blue).

Geometric alignment markedly improved cross-animal event detection (Fig. 3). When transferring the model from monkey T to monkey N, alignment reduced the movement-onset MAE from 370.45 ± 34.40 ms to 33.40 ± 7.12 ms and the turning-point MAE from 162.85 ± 27.78 ms to 48.64 ± 2.97 ms. Similarly, when transferring the model from monkey N to monkey T, alignment reduced the movement-onset MAE from 278.14 ± 20.65 ms to 27.50 ± 8.63 ms and the turning-point MAE from 113.80 ± 27.16 ms to 27.23 ± 5.54 ms. A complete summary of within-animal and cross-animal performance is provided in Table 1.

**Table 1.** Kinematic event-detection performance within and between animals. Values are the mean absolute error in milliseconds and are reported as the mean ± the standard deviation across 10 independent model runs. In the between-animal conditions, the model was trained on the animal other than the one identified in the first column. MAE, mean absolute error.

| Condition | Subject T |  | Subject N |  |
| --- | --- | --- | --- | --- |
|  | Movement Onset MAE | Turning Point MAE | Movement Onset MAE | Turning Point MAE |
| Within-subject training | 8.92 ± 2.03 | 23.93 ± 4.83 | 9.56 ± 3.64 | 39.31 ± 5.66 |
| Between -subject (unaligned) | 278.14 ± 20.65 | 113.80 ± 27.16 | 370.45 ± 34.40 | 162.85 ± 27.78 |
| Between -subject (aligned) | 27.50 ± 8.63 | 27.23 ± 5.54 | 33.40 ± 7.12 | 48.64 ± 2.97 |

**Figure 3.**
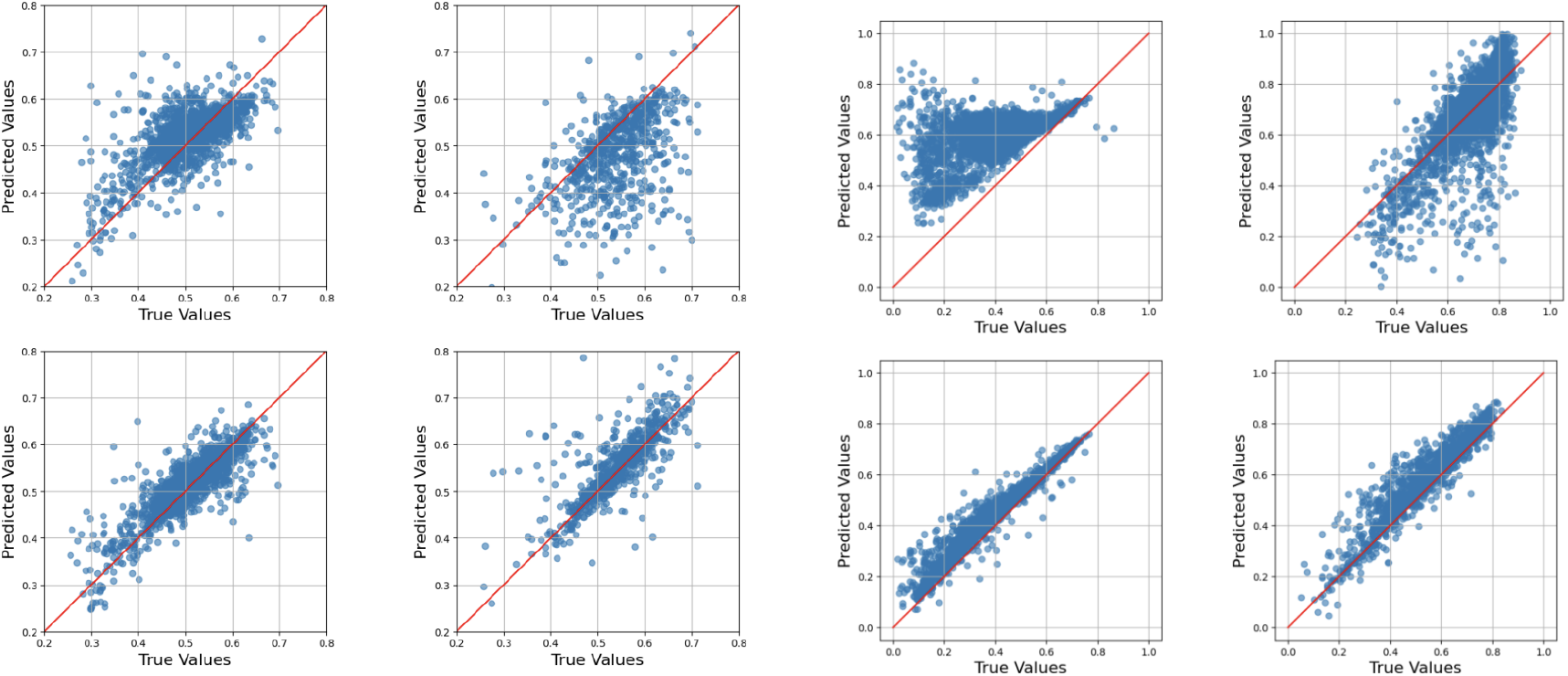
Inter-animal alignment improves kinematic event detection across animals. Predicted versus reference event times for turning-point detection (a) and movement-onset detection (b) before and after inter-animal alignment. In each panel, the top row shows predictions before alignment and the bottom row shows predictions after alignment. The left column shows models trained on monkey N and evaluated on monkey T, whereas the right column shows models trained on monkey T and evaluated on monkey N. The diagonal line represents perfect agreement between the predicted and the reference event times.

These findings indicate that geometric misalignment, rather than intrinsic differences in movement kinematics, accounted for a substantial fraction of the cross-animal prediction error. Following alignment, cross-animal performance approached within-animal levels, particularly for monkey T. The largest improvement was observed for movement-onset detection, suggesting that this event is especially sensitive to differences in coordinate-frame orientation. Turning-point detection also benefited from alignment, although to a lesser extent, consistent with the possibility that turning points depend more strongly on local trajectory features, such as changes in movement direction and velocity, that are less affected by global coordinate transformations.

### Blocking cerebellar output produces animal-specific changes in movement kinematics

Because the BiLSTM accurately detected kinematic events under control conditions, we next asked whether single-trial movement trajectories contained sufficient information to distinguish control reaches from those performed during reversible disruption of cerebellar output by high-frequency stimulation (HFS). Previous work has shown that this manipulation alters average motor behavior [19,20], but whether it can be identified from individual trials has not previously been established. To address this question, we used the same BiLSTM architecture to classify each trial as either control or HFS.

Within-animal decoding accuracy exceeded chance in both monkeys. In monkey T, the classifier achieved a mean accuracy of 71.0%, with target-specific accuracies ranging from 63.1% to 82.0%. In monkey N, the mean decoding accuracy was 61.9%, with target-specific accuracies ranging from 52.6% to 68.1% (Fig. 4). These results demonstrate that reversible cerebellar perturbation produces detectable alterations in the spatiotemporal structure of individual reaching movements. However, decoding performance differed substantially between animals and across movement directions, indicating that the behavioral consequences of cerebellar disruption are expressed in a subject- and task-dependent manner.

**Figure 4.**
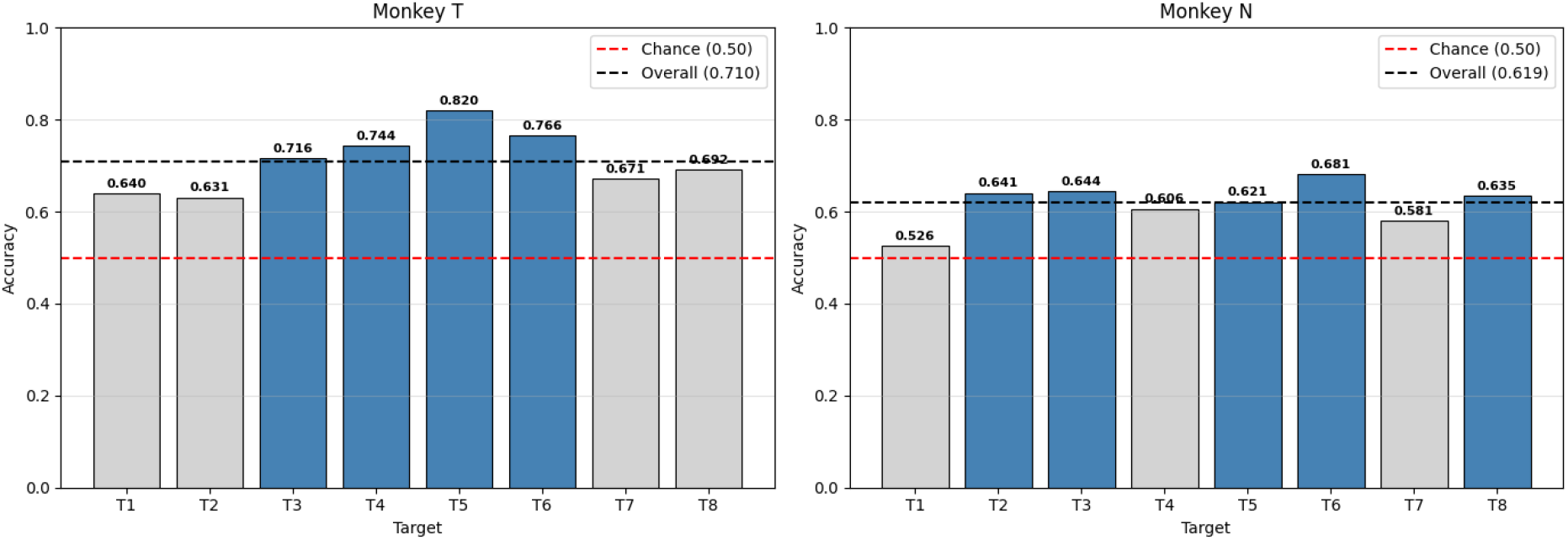
Within-animal decoding of HFS and control trials across reach targets. Bars show target-specific classification accuracy obtained from three-dimensional single-trial kinematics for **(a)** monkey T and **(b)** monkey N. The red dashed line indicates the nominal chance level of 50%, and the black dashed line indicates the overall decoding accuracy for each animal. Overall accuracy was 71.0% for monkey T and 61.9% for monkey N, with target-dependent variability in both animals.

We next examined whether the kinematic signatures of cerebellar perturbation generalized across animals. In contrast to the event-detection task, direct transfer of the BiLSTM classifier yielded decoding accuracies near the nominal chance level. A classifier trained on data from monkey T and tested on monkey N achieved an overall accuracy of 49.3%, whereas training on monkey N and testing on monkey T resulted in an accuracy of 47.7% (Fig. 5). Likewise, target-specific decoding accuracies remained close to chance, with no subset of movement directions showing reliable cross-animal generalization. These findings indicate that, unlike the kinematic features underlying event detection, the behavioral signatures of cerebellar perturbation were largely animal-specific and could not be captured by a model trained on another subject.

**Figure 5.**
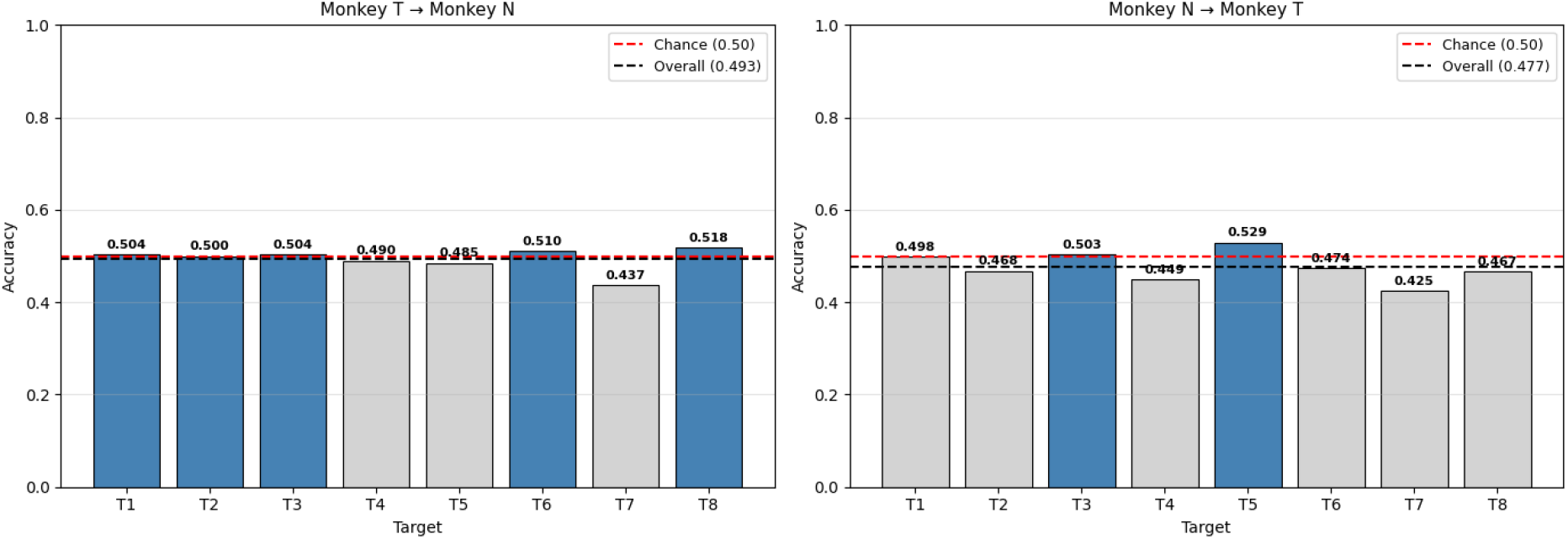
Inter-animal decoding of HFS and control trials across reach targets. Bars show target-specific classification accuracy obtained from three-dimensional single-trial kinematics for **(a)** a model trained on monkey T and evaluated on monkey N and **(b)** a model trained on monkey N and evaluated on monkey T. The red dashed line indicates the nominal chance level of 50%, and the black dashed line indicates the overall decoding accuracy for each transfer direction. Overall accuracy was 49.3% for T-to-N transfer and 47.7% for N-to-T transfer, with performance remaining close to chance across targets in both directions.

Applying inter-animal geometric alignment did not improve cross-animal decoding of cerebellar perturbation. In both transfer directions, decoding accuracy remained near the nominal chance level after alignment, indicating that the failure to generalize could not be explained primarily by differences in coordinate-frame orientation. This contrasts with the event-detection task, where geometric alignment was sufficient to recover near within-animal performance. Thus, whereas the kinematic features underlying movement-onset and turning-point detection were largely shared across animals after geometric normalization, the trajectory features distinguishing control and cerebellar perturbation trials remained animal-specific. These findings suggest that cerebellar disruption alters movement kinematics through subject-specific changes in movement dynamics rather than through a common geometric transformation of reaching trajectories.

## Discussion

This study demonstrates that a BiLSTM-based sequence model can accurately extract behaviorally relevant kinematic information from markerless tracking of unconstrained three-dimensional reaching movements on a single-trial basis. A principal finding is that, after correcting for differences in recording geometry, the network generalized across animals without requiring subject-specific retraining. Specifically, the model accurately detected movement onset and corrective turning points within individual animals, and this performance was largely preserved when applied to a second animal following geometric alignment. We further showed that the same network architecture could decode the behavioral consequences of reversible cerebellar perturbation from single-trial movement trajectories. In contrast to event detection, however, the kinematic signatures associated with cerebellar disruption did not generalize across animals and were not recovered by geometric alignment, indicating that they reflect subject-specific modifications of movement dynamics rather than differences in recording geometry.

These findings have three important implications. First, they establish an automated and accurate framework for detecting key kinematic events from unconstrained primate movements. Second, they demonstrate that a major source of variability limiting cross-animal generalization arises from differences in coordinate-frame geometry and can be effectively removed using a simple alignment procedure. Third, they show that, although single-trial movement kinematics contain sufficient information to decode experimentally induced changes in neural function, the behavioral expression of these perturbations remains highly individual. Together, these results establish a robust and broadly applicable framework for quantitative analysis of primate motor behavior while highlighting the distinction between invariant kinematic representations that generalize across subjects and individualized movement strategies that do not.

Within animals, the BiLSTM detected movement onset with high temporal precision, achieving mean absolute errors of less than 10 ms in both monkeys and consistently outperforming a conventional velocity-threshold approach [9,10]. This finding highlights the advantages of sequence-based models for extracting kinematic events from noisy and variable movement trajectories. Traditional threshold-based and statistical onset detectors have been widely used in motor behavior analysis [9–14], but their performance is often limited by sensitivity to signal quality, baseline fluctuations, threshold selection, and subject-specific parameter optimization. In contrast, the BiLSTM learns the temporal structure of the movement directly from the data, integrating information over the entire trajectory to estimate event timing from both preceding and subsequent kinematic context. This ability likely underlies its greater robustness to trial-to-trial variability and measurement noise.

Turning-point detection was consistently less precise than movement-onset detection. This difference likely reflects the intrinsic characteristics of the two events rather than a limitation of the model. Movement onset corresponds to a relatively abrupt transition from quiescence to sustained motion, producing a well-defined temporal landmark. In contrast, corrective turning points arise from a gradual reorganization of movement direction and velocity, the timing of which varies with the geometry of the target jump and the dynamics of the ongoing reach [21]. Consequently, turning points are expected to exhibit greater behavioral variability and less sharply defined temporal boundaries, making both manual annotation and automated detection inherently more challenging.

Despite the high within-animal performance, the model initially generalized poorly across animals. A straightforward solution would have been to retrain or fine-tune the network for each subject, but such an approach is labor-intensive, reduces reproducibility, and limits the practical utility of a generalizable event detector. Instead, our analysis revealed that a major source of the domain shift arose from differences in the geometric coordinate frames used to reconstruct the trajectories. In markerless three-dimensional tracking systems, the reconstructed coordinates are defined by camera placement and calibration, such that identical reaching movements recorded in different experimental configurations may appear arbitrarily rotated relative to one another.

Consistent with this interpretation, direct transfer of a detector trained on one animal to the other resulted in large localization errors despite the animals performing the same behavioral task. Correcting this geometric mismatch using orthogonal Procrustes alignment largely restored cross-animal performance, substantially improving detection of both movement onset and corrective turning points and reducing the gap between within-animal and cross-animal accuracy. These findings indicate that much of the apparent failure to generalize reflected differences in the representation of the trajectories rather than differences in the underlying movement kinematics. More broadly, they demonstrate that simple geometric normalization can effectively remove an important source of variability in markerless motion capture, enabling trained models to be transferred across subjects without additional retraining.

However, alignment did not fully restore performance to within-animal levels. This suggests that although geometric correction can resolve coordinate-frame discrepancies, other factors, such as variations in individual movement execution or limb anatomy persist and are likely to contribute to the remaining performance gap. This result has practical implications for studies that combine behavioral recordings across animals, sessions, or experimental systems. Cross-subject degradation may be misinterpreted as evidence that the learned behavioral representation is inherently subject-specific, when part of the apparent domain shift may in fact arise from arbitrary properties of the recording coordinate system. Correcting this geometric component before model transfer could reduce the amount of animal-specific annotation and retraining required for large behavioral datasets. At the same time, the remaining performance disparity indicates that rigid alignment should be treated as an initial harmonization step rather than as a complete solution to inter-animal variability.

The second objective of this study was to determine whether single-trial movement trajectories contain sufficient information to identify alterations in motor behavior induced by a reversible perturbation of the motor system. To address this question, we used data recorded during a transient blockade of cerebellar output, a manipulation known to impair reaching performance [19,20], and tested whether the BiLSTM could classify individual trials as either control or cerebellar block. In contrast to the event-detection task, the decoding analysis revealed substantial variability across animals and movement directions. Within each animal, decoding accuracy exceeded the nominal chance level, demonstrating that cerebellar perturbation produced detectable changes in the spatiotemporal organization of individual reaching trajectories. However, decoding performance was consistently higher in monkey T than in monkey N and varied across target directions, indicating that the behavioral consequences of cerebellar disruption depended on both the individual and the movement being performed. This directional dependence suggests that cerebellar perturbation does not simply produce a uniform reduction in movement speed or amplitude, but instead alters specific aspects of movement coordination in a context-dependent manner. Such an outcome is consistent with the established role of the cerebellum in coordinating multijoint movements and compensating for limb dynamics, including interaction torques and gravitational loads [22]. Because these biomechanical demands differ across reaching directions, the kinematic consequences of cerebellar disruption would also be expected to vary with target location, providing a plausible explanation for the observed target-dependent decoding performance.

Although within-animal decoding accuracy exceeded chance, decoding generalization across animals remained near the nominal chance level, even when the analysis was restricted to individual target directions. Unlike the event-detection task, geometric alignment failed to improve transfer performance. This contrast is particularly informative because the same alignment procedure successfully recovered much of the cross-animal performance for movement-onset and turning-point detection. Together, these findings suggest that the kinematic features supporting event detection are largely conserved across animals once differences in recording geometry are removed, whereas those distinguishing control from cerebellar perturbation are not.

A likely explanation is that event detection relies on fundamental (shared) kinematic transitions that are common to all reaches,, including the onset of movement and the local temporal structure surrounding corrective redirection. Once the coordinate frame was aligned, these features remained sufficiently similar to support transfer. However, decoding cerebellar perturbation depends on more subtle modifications of movement dynamics that reflect each animal’s response to the perturbation. Such differences could arise from individual compensatory strategies or differences in the physiological effects of the stimulation protocol used for blocking cerebellar outflow. Because the present study was not designed to distinguish among these possibilities, the relative contribution of each remains unresolved. Future work combining detailed behavioral analyses with direct physiological measurements of perturbation efficacy will be necessary to determine why the kinematic signatures of cerebellar disruption remain subject-specific despite successful geometric normalization.

Several limitations of this study should be acknowledged. First, the experiments were conducted in only two animals, limiting the extent to which the observed patterns of cross-animal generalization can be extrapolated to a broader population. Future studies involving larger cohorts will be required to determine whether the benefits of geometric alignment extend across a wider range of workspaces, movement strategies, and recording configurations. Second, the alignment procedure was limited to a rigid rotational transformation derived from target-direction vectors. Consequently, it cannot compensate for other sources of inter-subject variability, including differences in movement scale, limb morphology, nonlinear workspace distortions, temporal dynamics, or more complex subject-specific trajectory characteristics. More flexible alignment approaches may therefore further improve cross-subject generalization. Finally, the alignment procedure relied on known correspondences between target locations in the two animals and on sufficient trajectory data to estimate representative target-direction vectors. Although these requirements were readily satisfied in the structured center-out reaching task used here, it remains unclear how well the approach will generalize to less constrained behaviors, tasks with different spatial organizations, or datasets lacking common task-related or anatomical landmarks. Extending the framework to such settings represents an important direction for future research.

Several directions for future work emerge from the present study. Validation in larger multi-animal datasets will be essential to establish the robustness and generalizability of the proposed framework. Such studies should incorporate stringent cross-subject and cross-dataset validation schemes and compare the rigid geometric alignment used here with more flexible nonlinear registration and representation-learning approaches. Domain-adaptation techniques may further improve transfer by disentangling task-invariant kinematic features from subject-specific movement patterns. In addition, combining detailed kinematic analyses with simultaneous neural recordings could help determine whether the limited cross-animal generalization of cerebellar perturbation decoding arises primarily from differences in the underlying neural perturbation, individual compensatory strategies, or other sources of biological variability. More broadly, our findings demonstrate that successful cross-subject generalization requires distinguishing between variability introduced by the measurement process and variability that reflects genuine biological differences. Explicit geometric normalization effectively removes an important source of measurement-related variability, enabling robust transfer of kinematic event detectors across animals. In contrast, decoding perturbation-induced changes in movement depends on individualized movement dynamics that cannot be captured by geometric alignment alone. Together, these results provide a practical framework for quantitative analysis of unconstrained primate behavior while emphasizing the need for computational approaches that jointly account for geometric and biological sources of variability.

## Methods

### Ethics statement

This study was conducted on two adult female monkeys (*Macaca fascicularis*, wt 5–8 kg). All primate care and surgical procedures were in accordance with the Hebrew University Guidelines for the Use and Care of Laboratory Animals in Research, supervised by the Institutional Committee for Animal Care and Use (ethical approval number MD-19-15835-4).

### Experimental setup and data collection

Two adult female monkeys (*Macaca fascicularis*, weight 5-8 kg) were trained to sit in a primate chair and perform center-out reaches with the right hand on a vertical touchscreen positioned approximately 25 cm in front of them. Each trial began with the monkey acquiring a central target. After a center-hold period of 0.45–0.60 s, a peripheral cue appeared at one of the eight possible target locations uniformly distributed around the circumference of a circle. The monkey was required to maintain the central hold during a variable delay period of 0.50-0.80 s, after which the disappearance of the central target served as the GO signal. Following the GO signal, the monkey had 1s to reach the cued peripheral target. Successful reaches were rewarded with a drop of applesauce. On 20% of the trials, the cued target switched location 0–200 ms after the GO signal, requiring the monkey to redirect its ongoing or planned reach to the new target location. These trials are referred to as ‘update trials’. After training, the monkeys were implanted with a square recording chamber positioned over the contralateral upper-limb area of the motor cortex under general anesthesia. A round chamber was also positioned using stereotactic measurements to allow access to the ipsilateral superior cerebellar peduncle (SCP) [23,24]. A postoperative MRI was performed to plan the trajectory of SCP electrode insertion. After recovery and re-training, a bipolar concentric stimulating electrode (Microprobes for Life Science Inc) was inserted through the round chamber along this trajectory. Electrode placement was guided by monitoring stimulation-evoked multi-unit responses in the contralateral primary motor cortex, and the electrode was secured at the site producing the appropriate cortical response [23,24]. During task performance, the superior cerebellar peduncle was blocked using high-frequency stimulation (HFS). This stimulation protocol consisted of a train of biphasic rectangular-pulse stimuli (where each phase was 200 µs) applied at 130 Hz throughout the set of cerebellar block trials. Each train was delivered at a fixed intensity of 150–250 μA.

Upper-limb kinematics were recorded at 120 Hz using six synchronized infrared motion-capture cameras controlled with Motive software (OptiTrack Inc.). Behavioral event times from the touchscreen task, implemented in MATLAB using Psychtoolbox, were synchronized with the motion-capture data using a data acquisition system (Ripple Inc.). Three-dimensional coordinates of right upper-limb landmarks, including the shoulder, elbow, wrist, and base and tip of the middle finger, were extracted offline using DeepLabCut, a deep-learning-based markerless pose-estimation framework trained on manually annotated video frames [4]. Hand velocity profiles were computed from the reconstructed trajectory using three-point differentiation. Movement onset was defined as the earliest time at which tangential hand speed exceeded 20% of the trial-specific peak speed. For update trials with discernible trajectory redirection, the turning point was estimated from the tangential velocity profile using a two-component minimum-jerk sub-movement model. Specifically, the observed velocity profile was fit as the sum of two minimum-jerk components, consistent with previous studies modeling reaching trajectories as linear combinations of minimum-jerk sub-movements [25,26]. The fitted components were sorted by onset time, and the redirection time was defined as the first sample at which the second component exceeded 20% of its own peak velocity. Trials without a resolvable second component were excluded from the turning point estimation. For both movement onset and turning point estimations, manual review and corrections were made to the annotated time points when necessary. In total, over multiple recording days, the dataset comprised 13,330 trials from monkey T, including 1,974 annotated update trials, and 10,204 trials from monkey N, including 656 annotated update trials. The same dataset was used for both kinematic event detection and HFS trial decoding. For event detection, the supervised labels were the movement onset and corrective turning point times. For HFS decoding, trials were labelled as either HFS or control, and class-balanced subsets were used for training and evaluation.

### Feature Extraction

For both kinematic event detection and HFS trial decoding on the 3D free-reach dataset, each trial was represented as a multivariate time series of seven kinematic features derived from the raw hand position recordings. The feature vector at each timestep consisted of the three components of the re-centered and rotated hand position, a normalized timestamp, the instantaneous speed (scalar magnitude of the 3D velocity vector), the angle subtended by three consecutive positions (a local curvature proxy), and the Euclidean distance from the current position to the trial endpoint. Trajectories were re-centered to the origin and rotated so that target 5 defined the reference direction, ensuring that the spatial representation was consistent in each animal’s dataset. Another contextual variable encoded the trajectory type; namely, one of five update angles (0°, 45°, 90°, 135°, 180°) indicating whether and how far the target jumped after the Go signal. This categorical variable was represented by a learned embedding that projected each of the five types into a two-dimensional continuous space and was concatenated with the temporal summary produced by the recurrent encoder at classification time. All feature dimensions were standardized independently using column-wise z-scoring: the mean and standard deviation were estimated from the training set and then applied to rescale both training and test sequences, thus preventing data leakage across partitions. For the between-animal experiments, normalization statistics were computed for each animal’s own training set and applied separately, so that no information about the target animal’s kinematics contaminated the normalization. Within each mini-batch, variable-length sequences were zero-padded to the length of the longest sequence in that batch. The padded sequences were then packed prior to LSTM processing, ensuring that the recurrent computation was only performed over valid (non-padded) timesteps and that no spurious gradient flowed from the appended zeros.

### Model Architecture

Both the event detector and the HFS classifier were based on the same BiLSTM encoder architecture, but the models were trained separately for the two tasks. This architecture was well-suited to both tasks because identifying kinematic events and stimulation-related effects require the integration of information across the full movement sequence, including both the preceding and the subsequent temporal context.

A schematic representation of the HFS-classification architecture is shown in Fig. 6. The BiLSTM processed the input sequence, and its hidden states were summarized using length-aware mean pooling to create a single 100-dimensional embedding (*h*) for each trial, to ignore padding noise. For event detection, a regression head was attached to the embedding. This head predicted the event time as the mean (*μ*) and the variance (*σ*^2^) of a Gaussian distribution. The network was trained by minimizing the Gaussian negative log-likelihood loss, thus allowing the model to estimate both the expected event time and trial-specific predictive variance. For HFS classification, the trial embedding *h* was passed to a single linear output unit. Training minimized the binary cross-entropy loss to distinguish HFS trials from control trials. We ensured class balance during training to guarantee that the classifier learned genuine kinematic differences rather than relying on label frequency.

**Figure 6.**
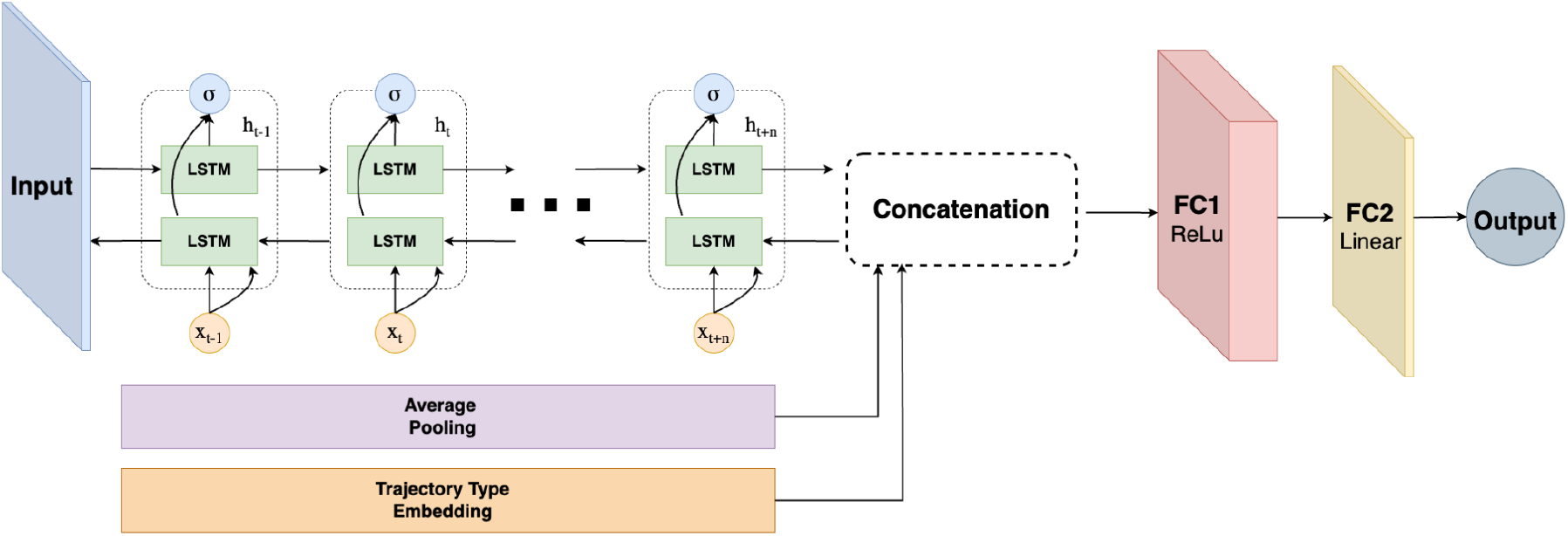
Schematic representation of the BiLSTM-based architecture used for HFS decoding. The input trajectory was represented as a multivariate time series of kinematic features and processed by a BiLSTM encoder. Hidden states over time were summarized by length-aware average pooling to generate a fixed-dimensional trajectory representation. For the HFS decoding task, this representation was concatenated with a learned embedding of trajectory type and passed through two fully connected layers to produce the output classification score.

### Training Protocol and Evaluation

All models were implemented in PyTorch and trained on CPU. Each dataset was partitioned into 80% training and 20% test using stratified sampling, with stratification performed jointly over trial label and target identity so that the per-target HFS/control balance was preserved in both subsets. The event detectors were trained for 100 epochs. All models used the Adam optimizer with a learning rate of 0.001 and a batch size of 32. To obtain stable performance estimates, all experiments were repeated over 10 independent runs using sequential random seeds (42–51), and the results are reported as the mean ± standard deviation across runs. Event detection performance was quantified by the mean absolute error (MAE) in milliseconds after inverting the normalization; HFS decoding performance is reported as classification accuracy on the balanced test set, overall and broken down by target identity.

### Inter-Animal Alignment (IAA)

Markerless 3D pose estimation systems, such as DeepLabCut, recover hand positions within a camera-defined world coordinate frame determined by multi-camera extrinsic calibration. This frame is arbitrary up to a rigid transformation across different experimental setups and animals. Consequently, trajectories directed at identical targets may appear rotated relative to one another, preventing the direct transfer of trained event detection models between subjects because the underlying kinematic features are not rotation-invariant. To resolve this geometric misalignment, we employed a rotation-constrained Procrustes alignment based on the Kabsch algorithm [18] to estimate the optimal rotation matrix that mapped one animal’s workspace onto the other. For each subject *i* and reaching target *j*, we determined the average final hand position relative to the workspace origin and normalized this vector to unit length. This produced a canonical target-direction vector *v*_*i*_,□ in three-dimensional space. To find the optimal rotation matrix *T\** belonging to the special orthogonal group SO(3) across *k* matching targets, we formulated the following optimization problem:

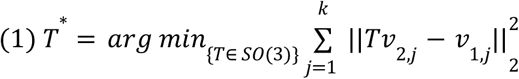

The cross-covariance matrix *M* was then constructed as:

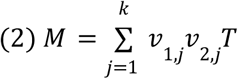

We performed a singular value decomposition on this matrix, such that *M = UΣ − V*^*T*^. The rotation matrix was subsequently calculated as:

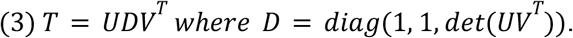

The inclusion of the correction matrix *D* ensured that the determinant of *T\** equaled +1, thus preventing improper rotations or reflections. This transformation was applied to align the source-subject trajectories with the target-subject coordinate frame before performing between-animal performance evaluations. This Inter-Animal Alignment (IAA) procedure placed both animals’ data in a shared geometric reference frame, thus correcting the coordinate-frame discrepancy introduced during 3D reconstruction while preserving all underlying relative distances and directions of movement.

## Acknowledgements

This work was funded by the Israel Science Foundation (ISF 1207/23, YP), the Deutsche Forschungsgemeinschaft (DFG, German Research Foundation Project-ID 431549029–SFB 1451) to YP, The Binational Science Foundation (BSF-2023321) for YP, the Council for Higher Education PhD Sandwich Fellowship, the Government of Israel to NS for undertaking his dual-PhD at the Hebrew University and Northwestern University, and the generous support of the Baruch Foundation to YP. The authors thank Ora Ben Harosh for her assistance with the training and the experiments involving the non-human primates used in this study. The authors also thank Dr. Ran Harel for assistance with the surgical procedures.

## Author contributions

J.G. and Y.P. conceived the study. N.S. performed experiments and preprocessed the data. A.M. analyzed data. A.M., N.S., Y.P. and J.G. wrote the manuscript. All authors reviewed and approved the final manuscript.

## Data availability

A subset of the data analyzed in this study is publicly available in the GitHub repository at https://github.com/TaliVas/monkey-alignment-network. The complete dataset is available from the corresponding author upon reasonable request.

## Code availability

The code used for data preprocessing, feature extraction, inter-animal alignment, model training, and evaluation is publicly available at https://github.com/TaliVas/monkey-alignment-network under the GNU General Public License v3.0. The repository also contains example data subsets and instructions for reproducing the analyses reported in this study.

## Additional information

The authors declare no competing interests. This study was performed on two adult female monkeys (*Macaca fascicularis*, weight 5-8 kg). All care and surgical procedures were in accordance with the Hebrew University Guidelines for the Use and Care of Laboratory Animals in Research, supervised by the Institutional Committee for Animal Care and Use (ethics approval number MD-19-15835-4).

## Notes

https://github.com/TaliVas/monkey-alignment-network

### Competing Interest Statement

The authors have declared no competing interest.

